# Efficacy of a novel SARS-CoV-2 detection kit without RNA extraction and purification

**DOI:** 10.1101/2020.05.27.120410

**Authors:** Tatsuya Fukumoto, Sumio Iwasaki, Shinichi Fujisawa, Kasumi Hayasaka, Kaori Sato, Satoshi Oguri, Keisuke Taki, Sho Nakakubo, Keisuke Kamada, Yu Yamashita, Satoshi Konno, Mutsumi Nishida, Junichi Sugita, Takanori Teshima

## Abstract

Rapid detection of SARS-CoV-2 is critical for the diagnosis of coronavirus disease 2019 (COVID-19) and preventing the spread of the virus. A novel “2019 Novel Coronavirus Detection Kit (nCoV-DK)” halves detection time by eliminating the steps of RNA extraction and purification. We evaluated concordance between the nCoV-DK and direct PCR. The virus was detected in 53/71 fresh samples by the direct method and 55/71 corresponding frozen samples by the nCoV-DK. The overall concordance rate of the virus detection between the two methods was 94.4% (95% CI, 86.2-98.4). Concordance rates were 95.2% (95% CI, 83.8-99.4), 95.5% (95% CI, 77.2-99.9), 85.7% (95% CI, 42.1-99.6) in nasopharyngeal swab, saliva, and sputum samples, respectively. These results indicate that the nCoV-DK effectively detects SARS-CoV-2 in all types of the samples including saliva, while reducing time required for detection, labor, and risk of human error.

## Introduction

Rapid and accurate detection of SARS-CoV-2 is critical for the prevention of outbreaks of coronavirus disease 2019 (COVID-19) in communities and hospitals. The diagnosis of COVID-19 is made by real-time quantitative PCR (RT-qPCR) testing of specimens collected by nasopharyngeal or pharyngeal swabs, with the nasopharyngeal route being the standard and sensitivity to the virus ranging from 52% to 71%(1-5). However, swab sample collection poses a risk of viral transmission to healthcare workers. Self-collecting saliva specimens are noninvasive tool for the virus detection and reduce a risk of health care workers. A series of recent studies have shown efficacy of saliva as a diagnostic tool(6-10). We recently reported 97% concordance rate between nasopharyngeal swab samples and saliva in the detection of SARS-CoV-2(11).

The 2019 Novel Coronavirus Detection Kit (nCoV-DK, Shimadzu Corporation, Kyoto, Japan) is a novel SARS-CoV-2 detection kit, which eliminates the steps of RNA extraction and purification by using the Ampdirect™ technology(12), thus significantly reducing the time required for sample preparation and PCR detection from more than 2 hours to about 1 hour. In addition, the risk of human error can be reduced by omitting the manual RNA extraction. However, nCoV-DK was initially developed for the use of nasopharyngeal swab specimens and it remained to be elucidated whether saliva samples could be applied to the nCoV-DK, since saliva has high RNase(13). In this study, we compared efficacy of the nCoV-DK with the direct PCR method requiring RNA extraction and purification using nasopharyngeal swab, saliva, and sputum samples.

## Methods

### Samples

Nasopharyngeal swab, sputum and saliva samples were collected from 9 patients who were admitted to our hospital after a diagnosis of COVID-19. A total of 71 samples were selected for this study according to the availability of the frozen stock samples. This study was approved by the Institutional Ethics Board and informed consent was obtained from all patients. Nasopharyngeal samples were obtained using FLOQSwabs (COPAN, Murrieta, CA, USA). Sputum and saliva samples were self-collected in a sterilized PP Screw cup 50 (Asiakizai Co., Ltd., Tokyo, Japan). 200 μL sputum or saliva were added to 600 μL PBS, mixed vigorously, then centrifuged at 20,000 X g for 5 minutes at 4°C, and and the supernatant was used.

### PCR

Direct RT-qPCR was performed according to the manual “Pathogen Detection 2019-nCoV Ver. 2.9.1” (March 19, 2020) from the National Institute of Infectious Diseases (https://www.niid.go.jp/niid/images/lab-manual/2019-nCoV20200319.pdf, accessed 2020-5-20). Total RNA was extracted using the QIAamp Viral RNA Mini Kit (QIAGEN, Hilden, Germany) from fresh samples. One-step RT-qPCR was performed using One-Step Real-Time RT-PCR Master Mixes (Thermo Fisher Scientific, Waltham, USA) and the StepOnePlus Real Time PCR System (Thermo Fisher Scientific) with forward primer (5’-AAA TTT TGG GGA CCA GGA AC-3), reverse primer (5’-TGG CAG CTG TGT AGG TCA AC-3’), and TaqMan probe (5’-FAM-ATG TCG CGC ATT GGC ATG GA-BHQ-3’).

The nCoV-DK PCR was carried out using the corresponding frozen specimens used for the direct PCR detection as above. The samples were processed according to the manufacturer’s instruction. In brief, 5 μL of sample and 5 μL of sample treatment reagent were mixed and heated for 5 min at 90°C using a thermal cycler to inactivate RNase. After cooling on ice, 15 μL of the reaction mixture containing primers and polymerase was added. PCR was performed using the CFX96 Touch Deep Well Real-Time PCR Detection System (Bio-Rad, California, USA). The nCoV-DK uses the “2019-nCoV_N1” and “2019-nCoV_N2” primer and probe sequences as described in the U.S. CDC’s “2019-Novel Coronavirus Real-time rRT-PCR Panel Primers and Probes “. N1 forward Primer (5’-GAC CCC AAA ATC AGC GAA AT-3’), N1 reverse Primer (5’-TCT GGT TAC TGC CAG TTG AAT CTG-3’) and N1 Probe (5’-ROX-ACC CCG CAT TAC GTT TGG TGG ACC-BHQ2-3’) were used. N2 Forward Primer (5’-TTA CAA ACA TTG GCC GCA AA-3’), N2 Reverse Primer (5’-GCG CGA CAT TCC GAA GAA-3’) and N2 Probe (5’-FAM-ACA ATT TGC CCC CAG CGC TTC AG-BHQ1-3’) were used. PCR positivity was defined as positive in either or both methods. Reagents and equipment for the nCoV-DK were provided by Shimadzu Corporation.

### Statistical analysis

Agreement between nasopharyngeal and saliva samples for the detection ability of SARS-CoV-2 was assessed using Cohen’s Kappa. Pearson’s correlation coefficient test was performed to identify the relation of the Ct values between the methods. All statistical analyses were performed with EZR (Jichi Medical University, Saitama, Japan), which is a graphic user interface for R (The R Foundation for Statistical Computing, Vienna, Austria). *P-value* of 0.05 was used as the cutoff for statistical significance.

## Results

Seventy-one specimens were tested by the direct PCR and the nCoV-DK. The virus was detected in 53 / 71 fresh samples by the direct PCR and 55 / 71 of the corresponding frozen samples by the nCoV-DK (Table 1). The overall concordance rate of the virus detection between the two methods was 94.4% (95% CI, 86.2-98.4). Interrater reliability of the two methods was strong (κ=0.85) determined by Cohen’s kappa analysis. Concordance rates were 95.2% (95% CI, 83.8-99.4), 95.5% (95% CI, 77.2-99.9), 85.7% (95% CI, 42.1-99.6) in nasopharyngeal swab, saliva, and sputum samples, respectively. Figure 1 shows a scatter plot presenting a comparison of Ct values in each positive sample between the two methods. There was a strong correlation between the two methods (r = 0.837, 95%CI = 0.736–0.902, P < 0.01). Significant correlations were also demonstrated in each sample type (Swab, r = 0.82, 95%CI = 0.673–0.905, P < 0.01; Saliva, r = 0.818, 95%CI = 0.507– 0.94, P < 0.01; Sputum, r = 0.945, 95%CI = 0.574–0.994, P < 0.01).

**Table 1.** Comparison of SARS-Cov-2 detection in the direct PCR and nCoV-DK method.

| nCoV-DK |  | Direct PCR |  | kappa (95%CI) |
| --- | --- | --- | --- | --- |
|  |  | Positive | Negative |  |
| Total | Positive | 52 | 3 | 0.85 (0.70-0.99) |
|  | Negative | 1 | 15 |  |
| Swab | Positive | 34 | 2 | 0.83 (0.60-1.00) |
|  | Negative | 0 | 6 |  |
| Saliva | Positive | 13 | 0 | 0.90 (0.72-1.00) |
|  | Negative | 1 | 8 |  |
| Sputum | Positive | 5 | 1 | 0.59 (0-1) |
|  | Negative | 0 | 1 |  |

**Figure 1.**
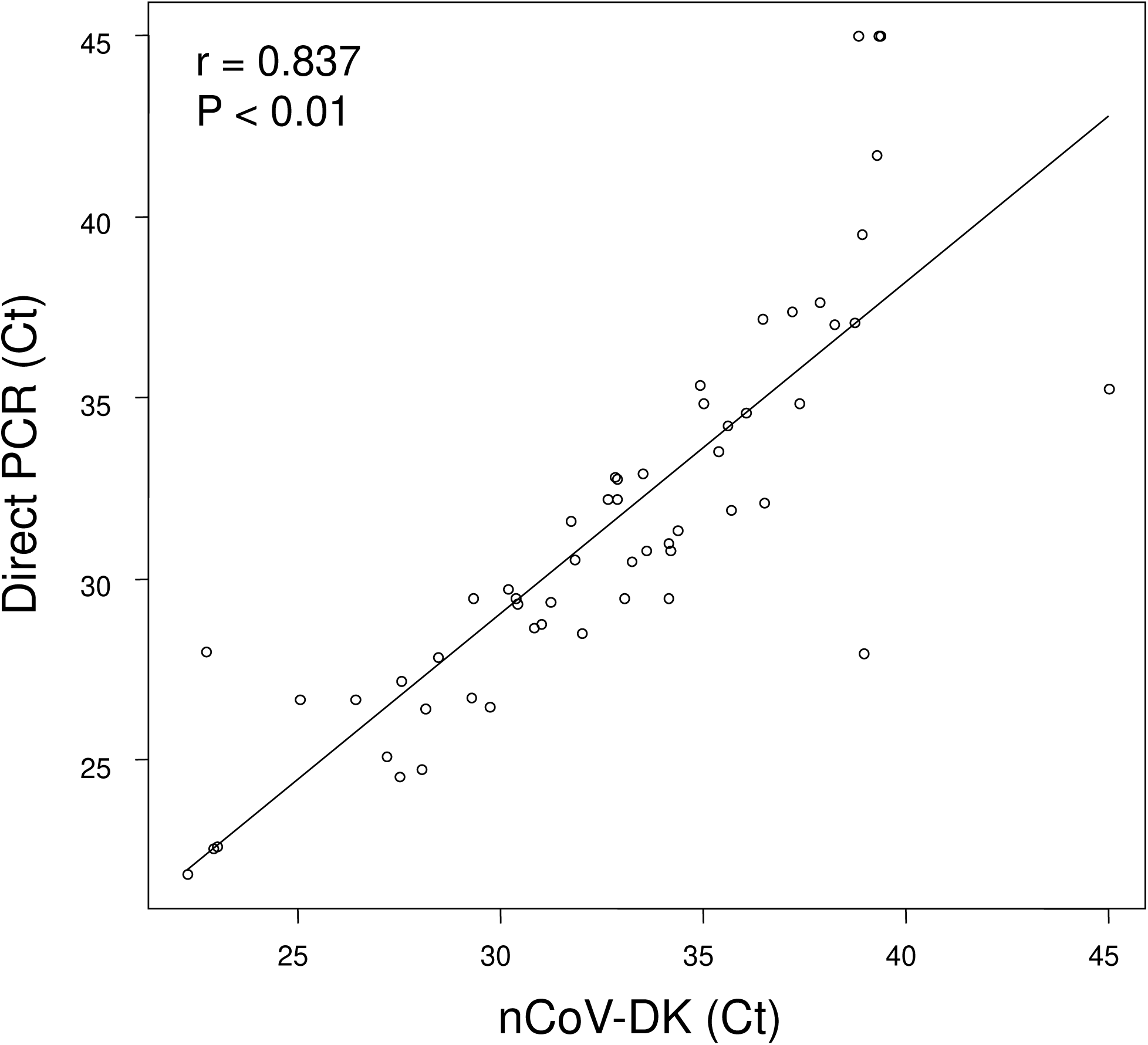
Correlation of Ct values between the direct PCR and nCoV-DK method. A scatter plot shows a comparison of Ct values between the two methods. Negative samples are denoted with a Ct of 45, which is the limit of detection.

## Discussion

In this study, we demonstrate that a novel SARS-CoV-2 detection kit nCoV-KD is as effective as the direct PCR methods in detecting SARS-CoV-2 in all types of the samples tested including saliva. Particularly, it should be noted that our study demonstrated that saliva is a reliable tool to detect the virus by the nCoV-KD even without process of RNA extraction and purification. However, there are some discordant results between the two methods. The virus was detected only by the direct PCR in one sample, while the virus was detected only by the nCoV-DK in 3 samples. It is unclear whether these are false-positive or true positive, since PCR primer sets are not the same between the two methods. In conclusion, the nCoV-DK has advantage over the direct PCR due to shorter detection time by eliminating the steps of RNA extraction and purification, without impairing diagnostic accuracy.

## Acknowledgements

This work was in part supported by grants from Japan Medical Association Research Institute.

